# Total, not isoform-specific, Lyn expression by macrophages promotes TLR activation and restricts proliferation

**DOI:** 10.1101/2025.07.07.663098

**Authors:** Anders J. Lindstedt, Joseph T. Greene, Tanya S. Freedman

## Abstract

Toll-like receptor (TLR) signaling is vital to antimicrobial macrophage function, and its dysregulation is associated with many disease states, including lupus, multiple sclerosis, pulmonary fibrosis, and cancer. The Src-family kinase Lyn plays activating and inhibitory roles downstream of TLRs, yet distinct functions of the Lyn splice variants LynA and LynB in TLR signaling had not been investigated. We used isoform-specific Lyn knockout mice (LynA^KO^ and LynB^KO^) to interrogate the contribution of each isoform to TLR signaling in bone marrow-derived macrophages. Bulk RNA sequencing and cytokine analyses revealed that complete Lyn deficiency (Lyn^KO^) dampens TLR4- and TLR7-induced inflammatory gene expression and TNF production, but enhances the expression of genes responsible for synthesizing the extracellular matrix and promoting proliferation. Despite a reduction in total Lyn levels, the expression of either LynA or LynB alone was sufficient to preserve wild-type transcriptional responses and TNF production in response to the TLR7 agonist R848. However, LyA^KO^ and LynB^KO^ macrophages did have partially impaired TNF production in response to the TLR4 agonist lipopolysaccharide. Additionally, LynA^KO^ and LynB^KO^ macrophages were hyperproliferative, like Lyn^KO^ cells. These data suggest that Lyn promotes macrophage activation downstream of TLRs and restrains aberrant proliferation and matrix deposition in a dose-dependent rather than isoform-specific manner.

**Summary Sentence:** RNA sequencing and functional assays demonstrate that both LynA and LynB restrict macrophage proliferation and drive TLR-induced ECM-remodeling and inflammatory cytokine production.

## Introduction

Macrophages play key roles in pathogen defense, wound healing, and tissue maintenance. Dysregulation of intracellular signaling is associated with infection (Campuzano et al., 2020), autoimmunity (Iwata et al., 2012; Scapini et al., 2010), fibrosis (Morse et al., 2019; Gibbons et al., 2011; Shen et al., 2014), and cancer progression (Jalil et al., 2020; Zhang et al., 2013; Barkal et al., 2019). Yet, mechanistic questions about how cells restrain pathological activation remain. Macrophage signaling can be initiated by transmembrane Toll-like receptors (TLRs), which detect extracellular ligands (e.g., TLR4) or endosomal ligands (e.g., TLR7) (Lin et al., 2017; Barton et al., 2006). TLRs respond to a variety of stimuli, including bacterial membrane components like Lipopolysaccharide (LPS), RNA and DNA motifs like GU-/AU-rich single-stranded RNA and unmethylated CpG DNA (Forsbach et al., 2008; Latz et al., 2004), and endogenous ligands like high mobility group box 1 and heat shock proteins (Park et al., 2004; Fang et al., 2011). Receptor ligation drives a diverse array of cellular responses. Inflammation results from the production of cytokines, such as tumor necrosis factor (TNF), interleukins (ILs), and interferons (IFNs). Chemokines, like C-C motif chemokine ligands (CCLs) and C-X-C motif chemokine ligands (CXCLs), are secreted, recruiting immune cells to the local environment (Kawai and Akira, 2011). TLRs also trigger cell proliferation via cyclin production (Li et al., 2010) and extracellular matrix (ECM) remodeling via matrix metalloprotease (MMP) and collagen or laminin synthesis (Lim et al., 2006; Bhattacharyya et al., 2013).

Signaling downstream of TLRs is established through interaction with the adaptor protein MYD88, although TLR4 also signals through the adaptor protein TRIF (Akira et al., 2004; Kawai and Akira, 2011). MYD88-dependent TLR4 signaling progress through MAPK and NF-ĸB pathways, culminating in the nuclear translocation of transcription factors NF-ĸB, CREB, and AP1-family members c-Jun and c-Fos (Duan et al., 2022), whereas TRIF-dependent signaling effectuates interferon regulatory factor 3 (IRF3) translocation (Duan et al., 2022). Similarly, TLR7 drives NF-ĸB and AP-1 nuclear translocation, however it also prompts IRF5 and IRF7 translocation (Wang et al., 2024). Even though these transcription factors regulate unique subsets of target genes, they converge on shared pathways. NF-ĸB induces inflammatory gene expression alone (e.g., *Il1b*) and in cooperation with IRF5 (e.g., *Tnf, Il6, Il12*) (Liu et al., 2025). AP-1 drives the expression of ECM-remodeling genes (e.g., *Mmp9)*, while also promoting *Tnf* and *Il6* transcription (Lee et al., 2009). CREB regulates macrophage survival through *Serpinb2, Bcl2, Il10*, and *Dusp* expression (Park et al., 2005; Sanin et al., 2015). IRF3 propels type I IFN responses through *Ifnb1* expression and triggers chemokine expression (e.g., *Cxcl10, Ccl5*) (Aziz et al., 2020), whereas IRF7 induces *Ifna1* expression in addition to *Ifnb1* (Lazear et al., 2013). Despite advances in our understanding of TLR signaling, the upstream regulatory factors that dictate selective activation and integration of these transcriptional programs remain incompletely defined.

The Src-family kinase (SFK) Lyn has emerged as a key upstream modulator of TLR signaling, but the breadth of TLR-induced transcriptional programs that are regulated by Lyn in macrophages remains underappreciated. Lyn serves both activating and inhibitory roles in TLR signaling (Brian et al., 2021; L’Estrange-Stranieri et al., 2024; Scapini et al., 2011). Global Lyn knockout (Lyn^KO^) mice develop a systemic lupus-like disease, characterized by myeloproliferation and splenomegaly, systemic inflammation, autoreactive antibodies, and glomerulonephritis (Hibbs et al., 1995; Chan et al., 1997; Brian et al., 2022). Progression to autoimmunity depends on the inflammatory environment created by IL-6, likely produced by hyperactive B cells (Tsantikos et al., 2010), and B cell-specific loss of Lyn is sufficient to drive the disease, including myeloproliferation and glomerulonephritis (Lamagna et al., 2014). Interestingly, dendritic cell (DC)-specific loss of Lyn is sufficient to drive disease, and this disease is dependent on the expression of MYD88 (Lamagna et al., 2013), as well as CARD9 (Ma et al., 2019). Lyn can inhibit TLR signaling in myeloid cells, including DCs (Lamagna et al., 2013; Ma et al., 2019; Dallari et al., 2017) and macrophages (Keck et al., 2010; Borzęcka-Solarz et al., 2017), with Lyn^KO^ cells producing more type I IFNs (e.g., IFNα, IFNβ), TNF, and IL-6 than wild-type (WT) cells. Mechanistically, Lyn mediates the phosphorylation of IRFs, leading to polyubiquitination and degradation, triggering suppression of type I IFN production (Ban et al., 2016; Tawaratsumida et al., 2022). However, this may be unique to classical DCs (cDCs), as Lyn^KO^ plasmacytoid DCs (pDCs) produce fewer inflammatory cytokines than WT (Dallari et al., 2017) and macrophage-specific loss of Lyn does not spur the development of autoinflammatory disease (Ma et al., 2019). Thus, the impact of Lyn on TLR-induced cellular responses may differ by cell type.

Several studies have highlighted the activating functions of Lyn. Surprisingly, overexpressing Lyn in mice also precipitates a lupus-like inflammatory disease (Hibbs et al., 2002), and antibody-secreting cells from human lupus patients have increased, not decreased, *LYN* expression (Chen et al., 2024). In myeloid cells, including macrophages, Lyn required to propel inflammatory signaling pathways (Shio et al., 2009; Lim et al., 2015). Specifically, TLR-driven inflammatory cytokine production is dependent on Lyn (Smolinska et al., 2008; Li et al., 2016; Toubiana et al., 2015; Avila et al., 2012). Given the multifunctional nature of Lyn in cell signaling and inflammatory disease, and the diverse signaling programs controlling TLR activation and cellular responses, the role of Lyn in macrophage TLR-signaling cascades requires further investigation.

*Lyn* RNA is alternatively spliced to produce two isoforms, LynA and LynB, which differ by in an insert in the N-terminal unique region of LynA (Yi et al., 1991). LynA is uniquely regulated through polyubiquitination and degradation (Freedman et al., 2015; Brian et al., 2019) and may be the dominant driver of mast-cell degranulation (Alvarez-Errico et al., 2010). Conversely, overexpressed LynB associates more with inhibitory signaling proteins (Alvarez-Errico et al., 2010). Our group has developed isoform-specific LynA^KO^ and LynB^KO^ mice and discovered that LynB^KO^ and female LynA^KO^ mice develop a lupus-like disease with age (Brian et al., 2022). Female LynA^KO^ and LynB^KO^ exhibited myeloproliferation and increased macrophage expression of CD11c, suggesting enhanced mobilization from the bone marrow (Lu et al., 2022). Interestingly, female LynA^KO^ macrophages expressed higher amounts of the activation marker CD80/86 relative to LynA^KO^ male and WT cells. Still, few studies have examined isoform-specific functions of Lyn in macrophages, and the role of LynA or LynB in TLR signaling is unknown.

To investigate specific functions of LynA or LynB in TLR responses, we performed RNA sequencing and cytokine analyses in single-isoform and total Lyn^KO^ macrophages treated with TLR4 or TLR7 agonists. While a complete loss of Lyn impaired TLR4- and TLR7-induced inflammatory gene expression and TNF production, the expression of either LynA or LynB was sufficient to preserve WT transcriptional responses and cytokine production. However, LynA^KO^ and LynB^KO^ macrophages did have partially impaired TNF production in response to TLR4 stimulation. Additionally, all Lyn-deficient macrophages were hyperproliferative, including isoform-specific KO cells. These data suggest that Lyn promotes macrophage activation downstream of TLRs and restrains aberrant proliferation in a dose-dependent, rather than isoform-specific, manner.

## Materials and Methods

### Mouse strains and housing

C57BL/6- derived LynA^KO^, LynB^KO^, and Lyn^KO^ mice have been described previously (Chan et al., 1997; Brian et al., 2022). LynA^KO^ and LynB^KO^ are hemizygous for the remaining isoform and express WT-levels of LynB or LynA, respectively (Brian et al., 2022). Animal use was compliant with University of Minnesota/American Association for Accreditation of Laboratory Animal Care and National Institutes of Health policy, under Animal Welfare Assurance number A3456-01 and Institutional Animal Care and Use Committee protocol number 2209-40372A. Mice were housed in a specific-pathogen-free facility under the supervision of a licensed Doctor of Veterinary Medicine and supporting veterinary staff. Breeding and experimental mice were genotyped via real-time polymerase chain reaction (Transnetyx, Memphis, TN). Genotyping was confirmed by immunoblotting, when appropriate.

### Generation of bone-marrow-derived macrophages (BMDMs)

BMDMs were generated as described previously (Brian et al., 2020; Freedman et al., 2015). Briefly, bone marrow was isolated from femora and tibiae of mice, treated in hypotonic solution to remove erythrocytes, and seeded in non-tissue-culture-treated polystyrene plates (CELLTREAT, Ayer, MA; Cat. 229653) and cultured at 37°C, 10% CO_2_ in “BMDM10” medium: Dulbecco’s modified Eagle’s medium (DMEM, Corning Mediatech, Manassas, VA; Cat. 10-017-CM) with final concentrations of 10% heat-inactivated fetal bovine serum (FBS, Omega Scientific, Tarzana, CA; Cat. FB-11), 1 mM sodium pyruvate (Corning Mediatech; Cat. 25-000-CI), 6 mM L-glutamine (Gibco, Grand Island, NY; Cat. 25030-081), 1% Penicillin-Streptomycin (179 and 172 µM, respectively) (Sigma-Aldrich, St. Louis, MO; Cat. P4333-100ML), and 5% CMG14-12 supernatant as a source of macrophage colony-stimulating factor (M-CSF). After 7 d culture with medium refreshment, BMDMs were harvested with enzyme-free cell dissociation buffer (Gibco, Grand Island, NY; Cat. 13150-016), washed with phosphate-buffered saline (PBS, Cytiva, Logan, UT; Cat. SH30256.01), and counted for replating.

### Treatment with TLR agonists

BMDMs were resuspended in BMDM10 without M-CSF (“DMEM10”), replated, and rested overnight at 37°C, 10% CO_2_. Cells were then treated 2 h (for RNA sequencing or qRT-PCR) or 24 h (for cytokine analysis) with DMEM10 alone (-) or with 2 ng/ml LPS from *S. Minnesota* R595 (List Biological Laboratories, Campbell, CA; Cat. 304) or 20 ng/ml R848 (InvivoGen, San Diego, CA; Cat. tlrl-r848-1). Cellculture supernatants were transferred to microcentrifuge tubes (Eppendorf, Hamburg, Germany; Cat. 0030123611) for cytokine analysis. For RNA sequencing, cells were washed in PBS and lysed in TRIzol (Thermo Fisher Scientific, Waltham, MA; Cat. 15596018). Samples were then stored at −80°C.

### RNA sequencing

RNA was isolated from BMDM TRIzol lysates via chloroform extraction followed by RNeasy Mini Kit (Qiagen, Hilden, Germany; Cat. 74104). Samples from four mice of each genotype were subjected to poly-A selection to isolate mRNA and then bulk, next-generation sequencing (Illumina NovaSeq 6000 platform, performed by Azenta Life Sciences, South Plainfield, NJ). Sequence reads (17.5-27 x 10^6^ per sample) were trimmed using *Trimmomatic v.0.36* and mapped to the ENSEMBL *Mus musculus* GRCm38 reference genome using *STAR aligner v.2.5.2b*. Unique hit counts were determined using *featureCounts* in the *Subread* package *v.1.5.2* for downstream analysis of differential gene expression.

### DESeq2 analysis

Genes were filtered in *R v.4.4.3* to retain only those with ≥10 counts in ≥3 of the 4 biological replicates within any genotype/treatment. Differential expression analysis was performed using the *DESeq2* package *v.1.46.0*, with samples grouped by genotype and treatment in the design formula (∼ Group). Variance-stabilizing transformation (VST) was applied to normalized counts for visualization and unsupervised clustering. Principal component analysis (PCA) was conducted on the 500 most variable genes across all samples using the *prcomp* function in the *stats* package of base *R*, and results were visualized using the *ggplot* function in the *ggplot2* package *v.3.5.2*, with samples colored by genotype and treatment. Differentially expressed genes (DEGs) were identified using the *results* function in *DESeq2*, and pairwise comparisons between genotypes within each treatment condition were performed. The *results* function in *DESeq2* uses the Wald test to calculate log_2_(fold-changes) and *p*-values and the Benjamini-Hochberg False Discovery Rate correction to calculate adjusted *p*-values. Genes were defined as differentially expressed if they met both a Benjamini–Hochberg adjusted *p*-value <0.05 and an absolute fold-change >1.5. *DESeq2* output was annotated using ENSEMBL gene IDs mapped to gene symbols using the *biomaRt* package *v.2.62.1*. To assess shared and condition-specific differential gene expression between genotypes, Venn diagrams were generated to visualize the overlap of DEGs between genotypes across untreated, LPS-treated, and R848-treated conditions. Venn diagrams were created using the *venn.diagram* and *draw.triple.venn* functions in the *VennDiagram* package *v.1.7.3*, and plots were rendered using the *grid.draw* function in the *grid* package of base *R*. VST-normalized gene expression was visualized using the *pheatmap* package *v.1.0.12*, with row-wise scaling, Euclidean clustering of genes, and a scaled color palette to represent relative expression levels. The total distribution of differential gene expression between genotypes was visualized with volcano plots generated using the *ggplot* function in *ggplot2*, with log₂(fold-change) on the x-axis and –log₁₀(adjusted *p*-value) on the y-axis. Threshold lines were included to denote significance cutoffs (adjusted *p*- value <0.05 and an absolute fold-change >1.5), and color-coding was applied to distinguish relative expression changes, with red indicating significantly increased expression, blue indicating significantly decreased expression, and all others in gray.

### Gene set enrichment analysis (GSEA)

The GSEA desktop application *v.4.4.0* (Broad Institute, Cambridge, MA) was used to evaluate pathway-level differences between genotypes at steady state (-) and after LPS or R848 treatment. VST-normalized gene-expression matrices (generated from *DESeq2*) were used as input, with genes ranked by signal-to-noise. Comparisons were made between genotypes within each treatment condition using phenotype-based permutation (*n* = 1000). Gene identifiers were mapped from ENSEMBL IDs to official gene symbols using the MSigDB v.2025.1 Mm.chip annotation file. Enrichment testing was performed using 29 hallmark gene-sets of interest or 16 curated ECM-related gene sets. Gene sets with <15 or >500 genes were excluded. Enrichment was weighted, and results were filtered and visualized using default GSEA settings. Significant gene-set enrichment was defined by a nominal *p*-value <0.1.

### Quantitative reverse transcription polymerase chain reaction (qRT-PCR)

RNA was converted into complementary DNA (cDNA) using a qScript cDNA Synthesis Kit (QuantaBio, Beverly, MA), according to manufacturer’s instructions. cDNA products were diluted 1:10 in ultrapure water and subjected in technical triplicates to qRT-PCR using QuantStudio 3 PCR (Thermo Fisher Scientific) with PerfeCTa SYBR Green SuperMix (Quantabio). The primer sequences for *Cyclophilin* were (5’ TGCAGGCAAAGACACCAATG 3’/ GTGCTCTCCACCTTCCGT) and for *Tnf* were (5’ CCTCTTCTCATTCCTGCTTGTG 3’/ TGGGCCATAGAACTGATGAGAG 3’). For each reaction, an equivalent amount of water in triplicate was substituted for cDNA and served as negative control. Ct values were normalized to the housekeeping gene *Cyclophilin*, and mRNA fold changes were calculated using the ΔΔCt method (Livak et al., 2001).

### Flow cytometry

BMDMs from each cell preparation were resuspended in flow-cytometry buffer comprising PBS, 2% heat-inactivated FBS, and 2 mM ethylenediaminetetraacetic acid (EDTA), and cells were stained for viability with Ghost Dye Red 780 (Tonbo Biosciences, San Diego, CA; Cat. 13-0865-T500). Cells were then blocked with Fc Shield, Clone 2.4G2 (Tonbo Biosciences; Cat. 70-0161-U500) and stained for TLR4 with BV650 anti-mouse CD284/MD-2 Complex, Clone MTS510 (BD Biosciences, Franklin Lakes, NJ; Cat. BDB740615) in flow-cytometry buffer. Cells were then washed and treated with Cytofix/Cytoperm (BD Biosciences; Cat. 554722). Cells were then washed with BD Perm/Wash buffer (BD Biosciences; Cat. 554723) and stained for TLR7 with PE anti-mouse CD287, Clone A94B10 (BioLegend, San Diego, CA; Cat. 160003). Flow cytometry was performed on a BD LSRFortessa or LSR-Fortessa X-20 cytometer, and data were analyzed using FlowJo software *v.10.9.0* (FlowJo, Ashland, OR).

### Cell proliferation assay

BMDMs were generated from 3 mice of each genotype and resuspended in DMEM10. PBS-diluted CellTrace Violet (CTV, Thermo Fisher Scientific, Waltham, MA; Cat. C34557) was added to cell suspensions. Cells were washed and resuspended in BMDM10 medium, plated in untreated polystyrene plates and cultured 96 h at 37°C in 10% CO_2_. Cells were then washed with flow-cytometry buffer, stained for viability, and analyzed via flow cytometry, as described above. The *Proliferation Modeling* function in FlowJo was used to quantify division within the “Live” cell gate, and the proportion of live cells within each generation was depicted graphically.

### Enzyme-linked immunosorbent assay (ELISA)

TNF secretion by BMDMs over 24 h was analyzed using the mouse TNF DuoSet ELISA Kit according to manufacturer’s instructions, with a 7-point standard curve (R&D Systems, Minneapolis, MN; Cat. DY410). A Tecan Infinite 200 PRO was used to determine the absorbance of each well at 450 nm (A_450_), with 540-nm background correction. The average zero standard was subtracted from the average of each standard or sample. A standard curve was created by plotting log(A_450_) by log[standard] and applying linear regression with GraphPad Prism *v.9.1.2* (GraphPad Software, Boston, MA). TNF levels were determined in reference to the A_450_ standard curve.

### Statistical analyses

Data presentation and statistics were performed using GraphPad Prism software. Data are presented as mean ± standard deviation. Significance was assessed via two-way ANOVA with Tukey’s multiple comparisons test. *p*-value <0.05 *, <0.01 **, <0.001 ***, <0.0001 ****, *ns* indicates no significant differences. Outlier analyses were performed on ELISA data using unbiased robust regression and outlier elimination (ROUT) with Q=1%. *n* indicates the number of biological replicates, where each replicate represents cells from an individual mouse. In graphs depicting proliferation or ELISA data, squares indicate cells derived from male mice, and circles indicate cells derived from female mice.

## Results

### Single-isoform expression of Lyn in macrophages is sufficient to maintain a WT-like transcriptome

We performed RNA sequencing on WT, LynA^KO^, LynB^KO^, and total Lyn^KO^ BMDMs following treatment with the TLR4 agonist LPS or the TLR7 agonist R848. PCA revealed that treatment with either LPS or R848 induced profound transcriptional changes that obscured the effects of Lyn expression. **(Fig. 1A)**. However, Lyn^KO^ BMDMs were shifted closer to steady-state transcriptomes than other genotypes. Many genes were differentially expressed (DEGs) according to treatment condition and genotype **(Fig. 1B)**. Even in the absence of TLR stimulation, Lyn^KO^ and WT BMDMs had >600 DEGs, suggesting that Lyn is highly involved in regulating the macrophage steady state **(Fig. 2A)**. Whereas the complete loss of Lyn led to significant upregulation/downregulation of many gene products **(Fig. 2B)**, the single-isoform knockouts had more modest, intermediate effects **(Fig. 1B)**. As expected, Lyn mRNA was significantly reduced in Lyn^KO^ BMDMs; moreover, TLR4 and TLR7 expression levels were not significantly different **(Fig. 2C)**. Lyn^KO^ BMDMs had reduced expression of genes encoding pro-inflammatory cytokines, such as *Tnf, Il1a,* and *Il1b,* and chemokines, such as *Ccl2, Ccl3, Ccl7,* and *Cxcl10*. Complete loss of Lyn also affected expression of genes coding for proteolytic enzymes and structural proteins, with decreased *Mmp8, Mmp12,* and *Mmp14* and increased *Col4a1, Col4a2,* and *Lama3*. Lyn^KO^ cells had increased expression of *Top2a, Tk1, Stmn1, Odc1, and Lig1*. These genes encode critical enzymes for DNA synthesis, replication, and repair, as well as cell-cycle progression and mitosis.

**Figure 1.**
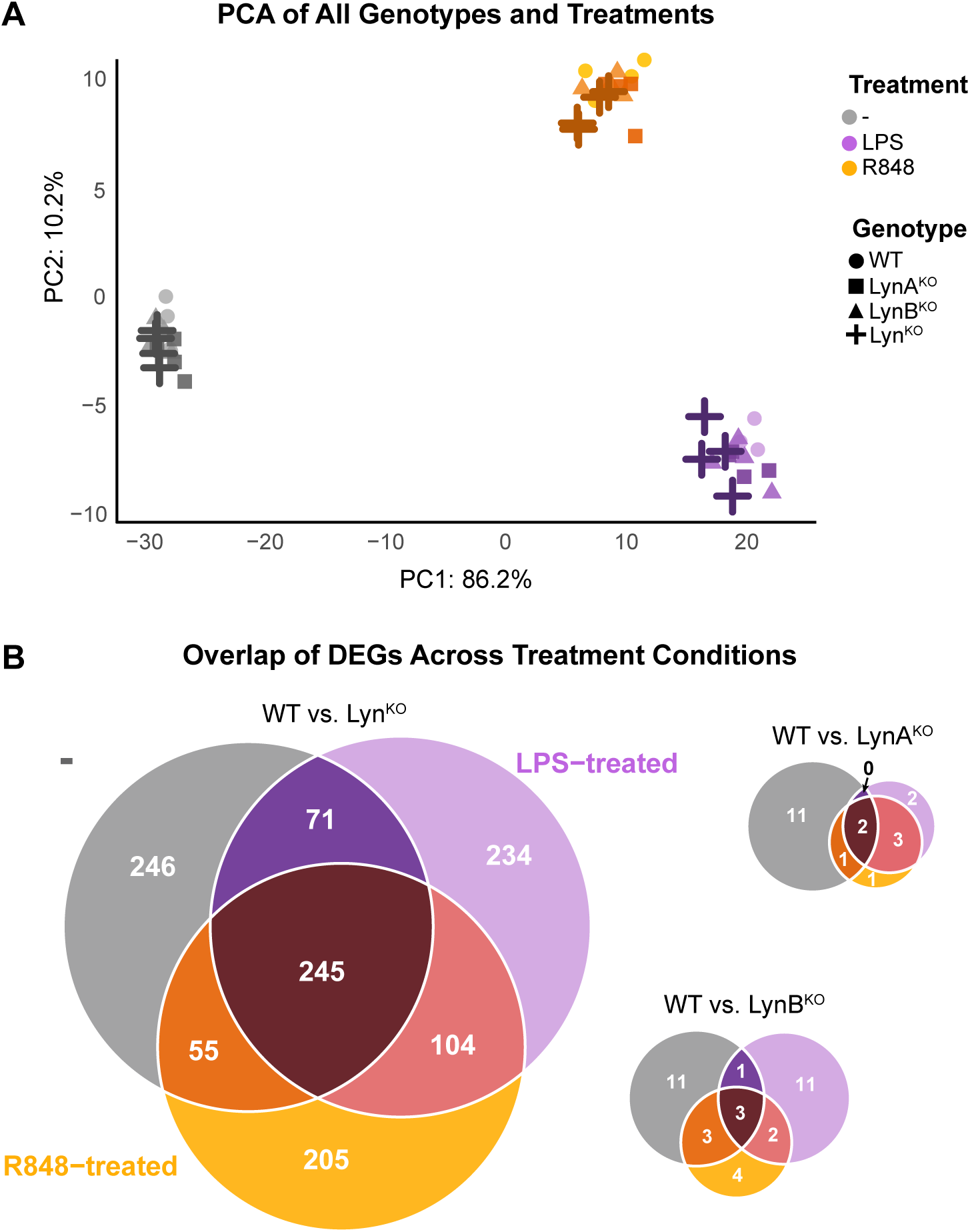
Lyn^KO^ BMDMs have altered steady-state and TLR-induced gene expression, whereas LynA^KO^ and LynB^KO^ transcriptomes resemble WT. **(A)** PCA of bulk RNA sequencing data from WT (circle), LynA^KO^ (square), LynB^KO^ (triangle) or total Lyn^KO^ (plus) BMDMs treated 2 h with medium alone (-, gray), 2 ng/ml LPS (purple), or 20 ng/ml R848 (orange) (*n*=4). Data set: 500 most variable genes, calculated from VST-normalized hit counts using *prcomp* in *R*. PC1 and PC2 account for 96.4% of the total variance. **(B)** Venn diagrams showing the overlap of significant DEGs across all treatments between Lyn knockouts and WT. DEGs were calculated from pairwise comparisons using *DESeq2* and defined by an absolute fold change >1.5 and an adjusted *p*-value <0.05. Biological replicates represent BMDMs from two male and two female mice of each genotype, independently differentiated and treated.

**Figure 2.**
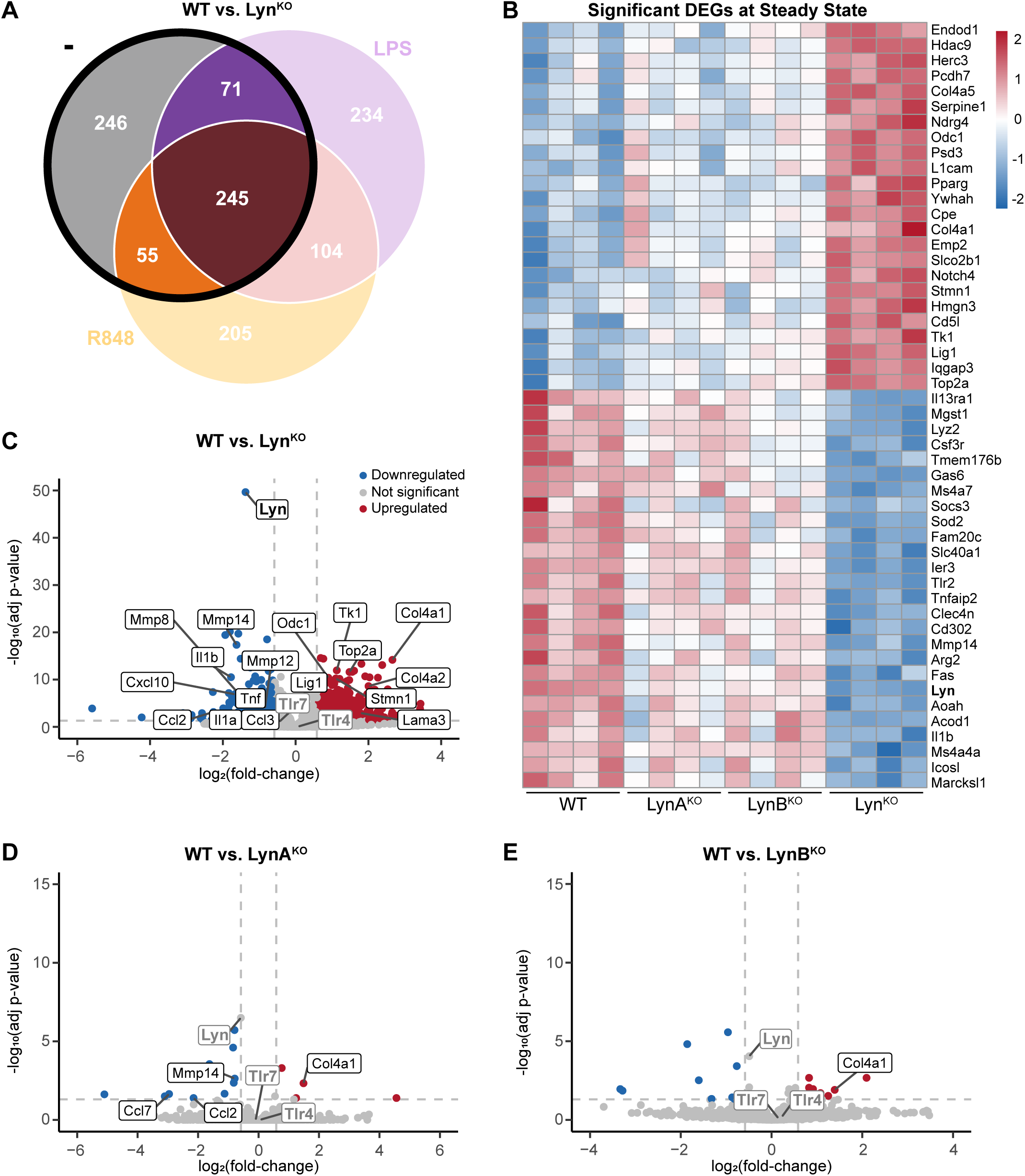
Lyn^KO^ BMDMs have decreased expression of genes encoding cytokines and proteases and increased expression genes encoding structural proteins and cell-cycle machinery. **(A)** Venn diagram highlighting DEGs in WT and Lyn^KO^ BMDMs at steady state (medium alone, -). **(B)** Heat map showing relative expression of the 50 highest-significance DEGs of WT and Lyn knockout BMDMs at steady state. Heat maps in this and other figures were generated using *pheatmap* in *R* to show z scores of VST-normalized hit counts for each sample relative to the mean count for each gene across all samples (red: increased, blue: decreased). The arrangement of rows was generated using hierarchical clustering by Euclidian distance. **(C-E)** Volcano plots highlighting DEGs in steady-state **(C)** Lyn^KO^, **(D)** LynA^KO^, or **(E)** LynB^KO^ BMDMs relative to WT. In this and other figures, DEGs were calculated from pairwise comparisons using *DESeq2* and defined by an absolute fold-change >1.5 and an adjusted *p*- value <0.05.

Although LynA and LynB are differentially regulated posttranscriptionally (Freedman et al., 2015; Brian et al., 2019) and contribute differentially to autoimmune disease and monocyte/macrophage phenotypes (Brian et al., 2022), steady-state LynA^KO^ and LynB^KO^ BMDMs had few transcriptomic differences relative to WT or each other **(Fig. 1B)**. There were few differences in the steady-state transcriptomes of WT and LynA^KO^ **(Fig. 2D)** or LynB^KO^ **(Fig. 2E)** BMDMs or between LynA^KO^ and LynB^KO^ **(Sup. Fig. 1A)**. However, LynA^KO^ cells had reduced expression of *Ccl2*, *Ccl7*, and *Mmp14*, and both LynA^KO^ **(Sup. Fig. 4B)** and LynB^KO^ **(Sup. Fig. 4G)** cells had increased expression of *Col4a1*. These findings suggest that, at steady state, Lyn^KO^ BMDMs already have transcriptomic changes that may alter their function and responses to stimuli. Moreover, expression of either Lyn isoform is sufficient to restore a WT-like transcriptome in resting cells.

To enable interpretation of subsequent analyses and confirm the observations that TLR4 and TLR7 levels were unaffected at the RNA level by Lyn knockout **(Sup. Fig. 2A)**, we assessed the effect of Lyn deficiency on the expression of TLR4 and TLR7 protein. Using flow cytometry, we assessed TLR4 and TLR7 surface protein in WT, LynA^KO^, LynB^KO^, and Lyn^KO^ BMDMs using flow cytometry **(Sup. Fig. 2B)**. As with mRNA, we found no differences in surface expression of the receptor proteins **(Sup. Fig. 2C)**.

### Few receptor-specific transcriptional differences distinguish TLR4 and TLR7 signaling in macrophages

We next assessed the most significant DEGs in WT BMDMs at steady state and after treatment with the TLR4 agonist LPS or the TLR7 agonist R848. For these studies, we chose agonist doses that induced comparable upregulation of *Tnf* in WT BMDMs **(Sup. Fig. 3A inset)**. Consistent with previous studies (Raza et al., 2014; O’Connell et al., 2017; Johannessen et al., 2018; Shi et al., 2021; Lin et al., 2017), treatment with either LPS or R848 drove upregulation of genes encoding pro-inflammatory cytokines (e.g., *Tnf, Il1a, Il6, Il12a, Il12b, Il23a, Acod1*), chemokines (e.g., *Ccl4, Ccl5, Cxcl1, Cxcl2, Cxcl3*), mitogens (e.g., *Csf2*), and matrix metalloproteases (e.g., *Mmp13*) (**Fig. 3A)**. Either TLR pathway also drove downregulation of *Cxcr4*, which, in vivo, leads to myeloid-cell egress from the bone marrow into peripheral blood (Kim et al., 2006). Focusing on transcriptomic difference uniquely modulated by the TLR4 or TLR7 pathway, LPS treatment uniquely drove interleukin and chemokine genes, such as *Il33* and *Cxcl9* **(Fig. 3B, Sup. Fig. 3B)**, and triggered a greater degree of gene induction than R848, with more upregulation of *Cxcl10* **(Sup. Fig. 3A)**. Macrophage-produced CXCL9 and CXCL10 are critical for anti-tumor T-cell infiltration and response to immune checkpoint blockade (House et al., 2020). Interestingly, R848 induced downregulation of several genes, including *Ankrd6*, *Mcc*, *Trim15*, and *Trim25* (**Fig. 3C, Sup. Fig. 3C)**. TRIM25 shifts the balance of signaling-pathway activation in macrophages, favoring MAPK and anti-inflammatory signaling over NF-ĸB activation (Liu et al., 2020). R848 also drove unique upregulation of genes like *Ifngr1*, *Il10ra*, and *Sirpa* (**Fig. 3C, Sup. Fig. 3C)**. A delicate balance of signaling through the IL10 receptor and SIRPα regulates inflammation-induced phagocytosis of healthy cells in macrophages (Bian et al., 2016). Despite some receptor-specific differences in gene induction, in WT BMDMs, most of the significant transcriptomic changes induced by TLR4 or TLR7 stimulation are shared between these two receptors.

**Figure 3.**
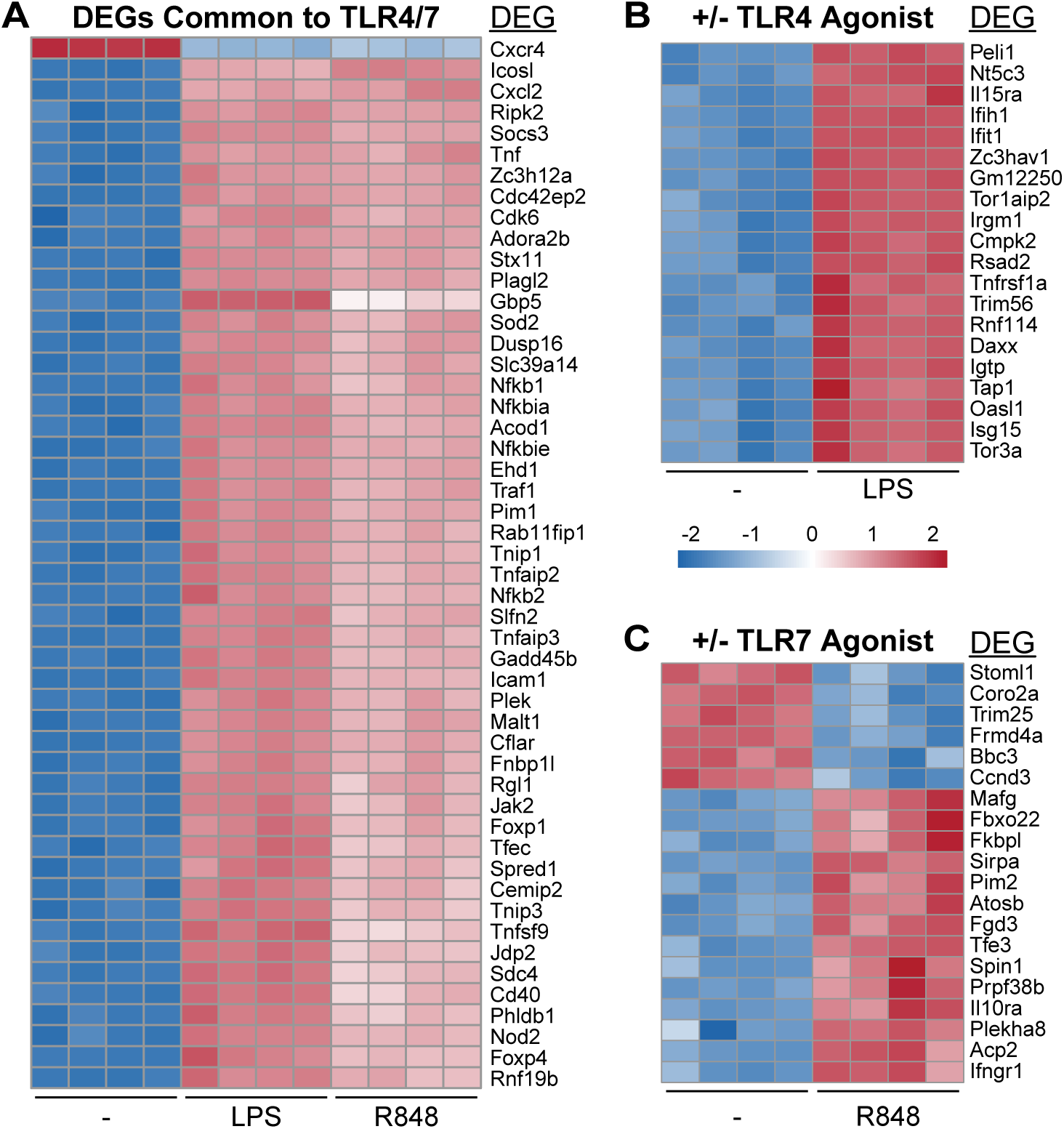
WT BMDM transcriptomes have some distinct features in LPS- and R848-treated conditions. **(A)** Heat map of the 50 most significant DEGs common to LPS-treated and R848-treated WT BMDMs compared to cells in medium alone. **(B,C)** Heat maps of the 20 most significant DEGs unique to **(B)** LPS treatment or **(C)** R848 treatment. Heat maps and DEGs were compiled as described in Fig. 2.

### Lyn deficiency broadly impacts TLR-induced gene transcription in macrophages

We then compared the transcriptomes of WT, LynA^KO^, LynB^KO^, and Lyn^KO^ BMDMs treated with TLR4 or TLR7 agonists. LPS or R848 treatment of Lyn^KO^ BMDMs lead to dysfunctional modulation of 371 genes that were also dysregulated at steady state (e.g., *Tnf, Il1a, Il1b, Ccl2, Ccl3, Cxcl10, Mmp8, Mmp12, Mmp14, Col4a1, Col4a2, Lama3*). However, Lyn^KO^ BMDMs failed to modulate the expression of 104 additional gene products after either TLR4 or TLR7 stimulation **(Fig. 4A)**, including failed upregulation of pro-inflammatory factors (e.g., *Il12b, Il23a, P2ry13, P2ry14, Pilrb1, Tnfsf15*) and chemokine-encoding genes (e.g., *Ccl22, Ccl24*), coupled with supraphysiological induction of inflammation-suppressing genes (e.g., *Traip, Sigirr)* (**Fig. 4B)**. Additionally, Lyn^KO^ cells had impaired induction of genes encoding matrix metalloproteases (e.g., *Mmp13*) and enhanced induction of genes encoding structural proteins (e.g., *Lama5, Plod2, Fgl2*). Again, isoform-specific LynA^KO^ **(Sup. Fig. 4A)** and LynB^KO^ **(Sup. Fig. 4F)** BMDMs displayed transcriptomes largely resembling WT cells. However, LynA^KO^ **(Sup. Fig. 4C)** and LynB^KO^ **(Sup. Fig. 4H)** BMDMs did have increased *Lama5* expression, and LynB^KO^ BMDMs had increased *Notch4* expression. Regardless, there were no significant transcriptional differences between LynA^KO^ and LynB^KO^ BMDMs after either LPS **(Sup. Fig. 1B)** or R848 **(Sup. Fig. 1C)** treatment.

**Figure 4.**
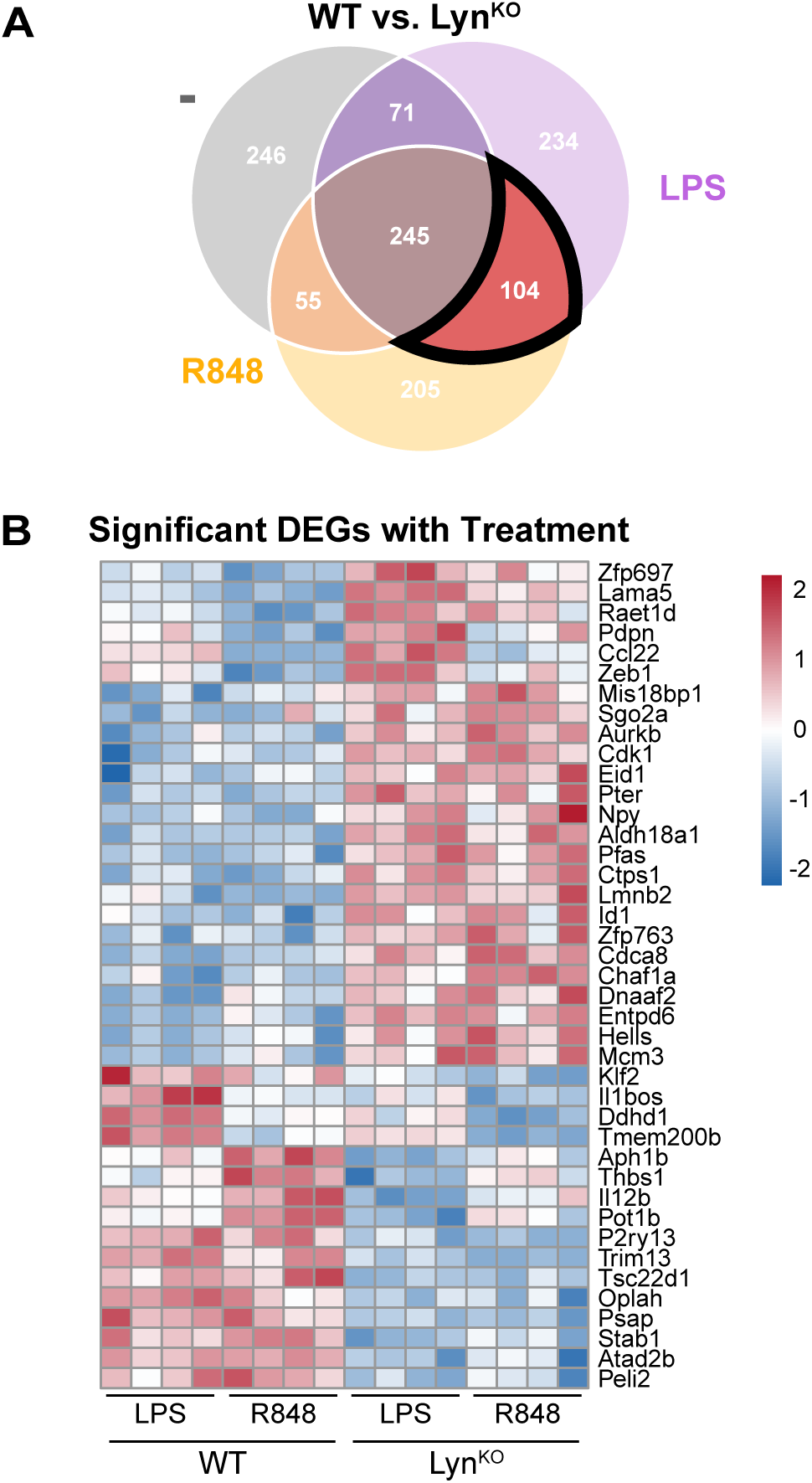
Lyn^KO^ BMDMs have impaired upregulation of a pro-inflammatory transcriptome after TLR stimulation. **(A)** Venn diagram defining the subset of genes differentially expressed after either LPS or R848 treatment of WT and Lyn^KO^ BMDMs. **(B)** Heat map showing the most significant DEGs in the subset defined in (A). Comparing the 50 most significant DEGs between Lyn^KO^ and WT BMDMs after LPS treatment with the 50 most significant DEGs after R848 treatment, 41 genes were common to both conditions. Heat maps and DEGs were compiled as described in Fig. 2.

To discern whether Lyn uniquely regulates signaling downstream of TLR4 or TLR7, we examined LPS-specific and R848-specific DEGs in WT and Lyn^KO^ BMDMs. We identified 234 DEGs found only in LPS-treated samples **(Fig. 5A)**. Gene products like *Jund* (an AP-1 family transcription factor), *Nupr1* (an autophagy suppressor), and *Pim1* (a Ser/Thr kinase that restricts cell growth) were uniquely downregulated, while *Traip* (an E3 ubiquitin ligase) and *Pkp3* (plakophilin, a component of desmosomes) were upregulated **(Fig. 5B)**. There were few LPS-specific DEGs between either LynA^KO^ **(Sup. Fig. 4D)** or LynB^KO^ **(Sup. Fig. 4I)** BMDMs and WT, but both genotypes had decreased expression of *Serpinb9*. In WT and Lyn^KO^ BMDMs, we identified 205 DEGs found only in R848-treated samples **(Fig. 5C)**. Gene products like *Tnfsf9* (4-1BBL, promotes T-cell co-stimulation) and *Mertk* (receptor tyrosine kinase) were uniquely downregulated, while *Jak3* (tyrosine kinase that mediates cytokine responses), *Jam2* (cellular-junction adhesion molecule), and *Timp1* (inhibits MMP activity) were upregulated **(Fig. 5D)**. There were no remarkable R848-specific DEGs between LynA^KO^ **(Sup. Fig. 4E)** or LynB^KO^ **(Sup. Fig. 4J)** BMDMs and WT. Although Lyn^KO^ BMDMs exhibited altered expression of some inflammatory signaling genes following either stimulus, the directionality and magnitude of these differences were not exclusive to one treatment, and no clear segregation of receptor-specific signaling pathways emerged. Most DEGs were not associated with canonical TLR signaling cascades, such as NF-κB, MAPK, or IRF-driven transcription. These data suggest that Lyn is required for full transcriptional activation downstream of both TLR4 and TLR7, and the absence of Lyn results in a broad attenuation of TLR-driven signaling rather than selective disruption of individual receptor-associated pathways.

**Figure 5.**
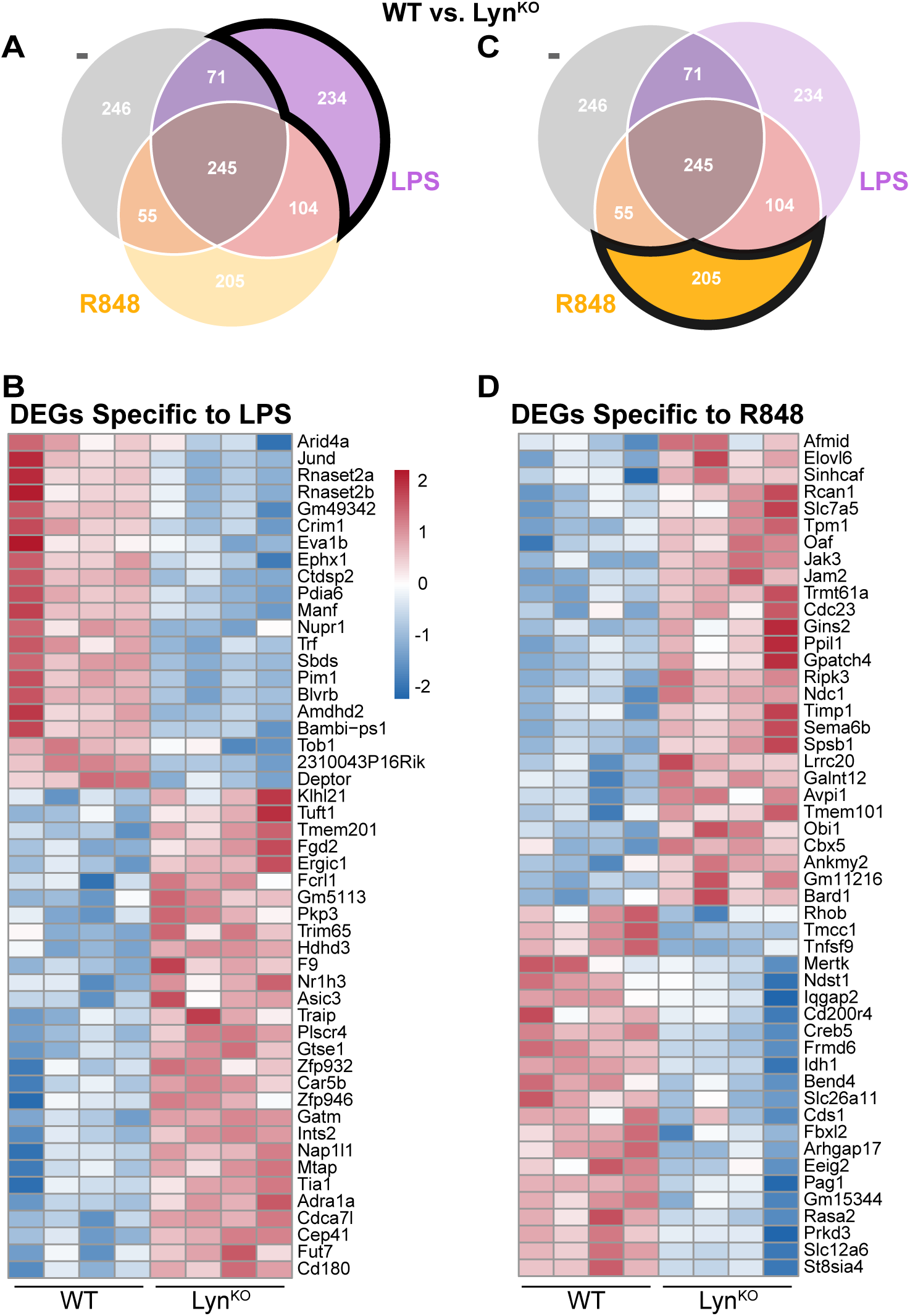
LPS- and R848-induced genes are differentially affected by Lyn knockout in BMDMs. **(A)** Venn diagram highlighting LPS-specific DEGs in WT and Lyn^KO^ BMDMs. **(B)** Heat map showing the 50 most significant LPS-specific DEGs in WT and Lyn^KO^ BMDMs. **(C)** Venn diagram highlighting R848-specific DEGs in WT and Lyn^KO^ BMDMs. **(D)** Heat map showing the 50 most significant R848-specific DEGs in WT and Lyn^KO^ BMDMs. Heat maps and DEGs were compiled as described in Fig. 2.

### Lyn restricts proliferation and promotes TLR-driven ECM remodeling and inflammatory responses

To refine our transcriptome-wide analyses of DEGs in WT and Lyn^KO^ BMDMs, we used gene-set enrichment analysis (GSEA) to determine which cellular functions appear to be most perturbed by the loss of Lyn **(Sup. Fig. 5A-C)**. We found basal enrichment of E2F-targeted **(Fig. 6A)** and mitotic-spindle-related **(Fig. 6B)** gene pathways in Lyn^KO^ BMDMs. As the E2F transcription factor and formation of a mitotic spindle are key components of cell proliferation (Liberzon et al., 2015; Johnson and Schneider-Broussard, 1998; Dang, 1999), we searched the DEG pool for other pro-mitotic gene products. Indeed, we found that Lyn^KO^, but not single-isoform knockout BMDMs, upregulate gene products promoting DNA synthesis, replication, and repair (e.g., *Tk1, Top2a, Lig1, Pcna, Mcm5*) and mitotic microtubule rearrangement (e.g., *Stmn1, Anln, Nusap1, Tpx2, Melk, Cit, Kif4, Spc25, Prc1, Ndc80, Plk1, Mad2l1, Espl1, Ncapd2*) (**Fig. 6C)**. To test the functional consequences of these transcriptional changes, we measured proliferation of WT, LynA^KO^, LynB^KO^, and Lyn^KO^ BMDMs in culture. Consistent with previous findings with Lyn^KO^ BMDMs (Scapini et al., 2009), we observed enhanced proliferation of Lyn^KO^ cells in culture, demonstrated by more CTV dilution in Lyn^KO^ BMDMs than WT **(Fig. 6D)**. Comparing parental and dividing cells at 96 h, we found that Lyn^KO^ BMDMs were significantly more likely to divide than WT **(Fig. 6E)**. Interestingly, though the transcriptional profile of LynA^KO^ and LynB^KO^ BMDMs only trended toward an intermediate phenotype, these cells also exhibited a greater degree of proliferation than WT in culture. This suggests that neither LynA nor LynB alone is sufficient to restrict cell proliferation, suggesting that a WT-like level of total Lyn is required to restrain cell division.

**Figure 6.**
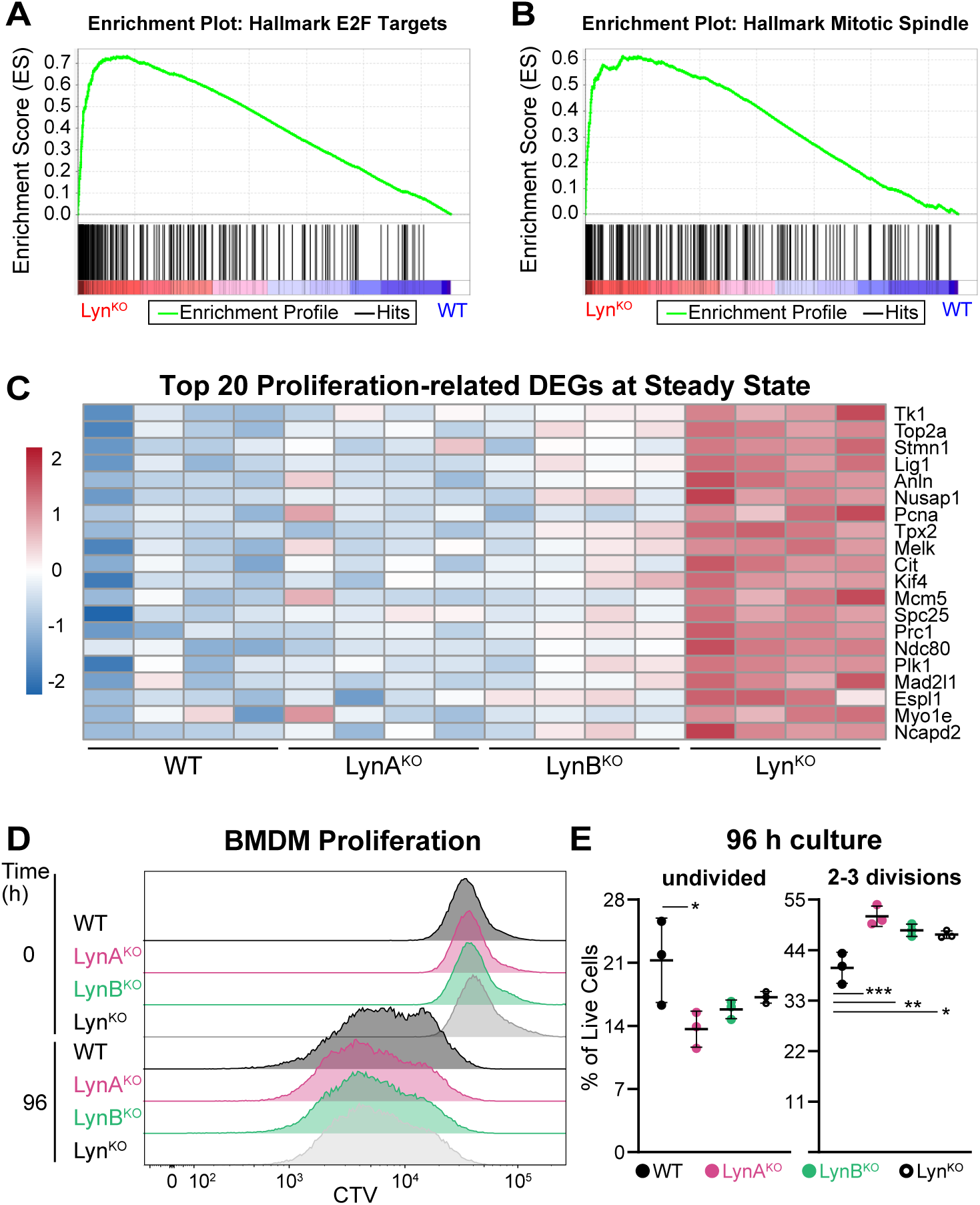
Lyn^KO^ BMDMs have enhanced cell proliferation at steady state. **(A,B)** GSEA showing enrichment of **(A)** E2F-targeted and **(B)** mitotic-spindle-related genes in Lyn^KO^ BMDMs at steady state. **(C)** Heat map of the 20 most significant DEGs related to proliferation in WT and Lyn^KO^ BMDMs. Heat maps and DEGs were compiled as described in Fig. 2. **(D)** Representative flow-cytometry histograms showing CTV fluorescence in BMDMs at steady state or after 96 h culture in M-CSF-containing medium. **(E)** Quantification of parental and dividing cells after 96 h (*n*=3). The parent generation was identified by the CTV peak at t=0, and subsequent generations were identified using FlowJo software. Data are presented as mean ± standard deviation. Significance was assessed via two-way ANOVA with Tukey’s multiple comparisons test. *p* values 0.01-0.03 *, 0.007 **, 0.001 ***. There were no significant differences between non-annotated pairs. *n*=3 biological replicates derived from different mice.

GSEA also revealed TLR-induced transcriptional changes in Lyn^KO^ BMDMs that favor ECM formation. After either LPS **(Fig. 7A)** or R848 **(Fig. 7B)** treatment, Lyn^KO^ cells had enhanced expression of core matrisome genes, with many of these having a greater magnitude of differential expression than at steady state. Notably, genes that prompt the synthesis of ECM components and expansion of the ECM (e.g., *Col4a1, Col4a2, Col4a5, Col4a6, Lama3, Lama5, Fgfr1, Fgf13, Pgf, Plod2*) were upregulated in Lyn^KO^ cells, while those that facilitate ECM degradation (e.g., *Mmp8, Mmp12, Adamtsl5, Slpi*) were downregulated relative to WT **(Fig. 7C)**. These data suggest that Lyn promotes ECM turnover, and defects in Lyn can lead to overgrowth of the ECM.

**Figure 7.**
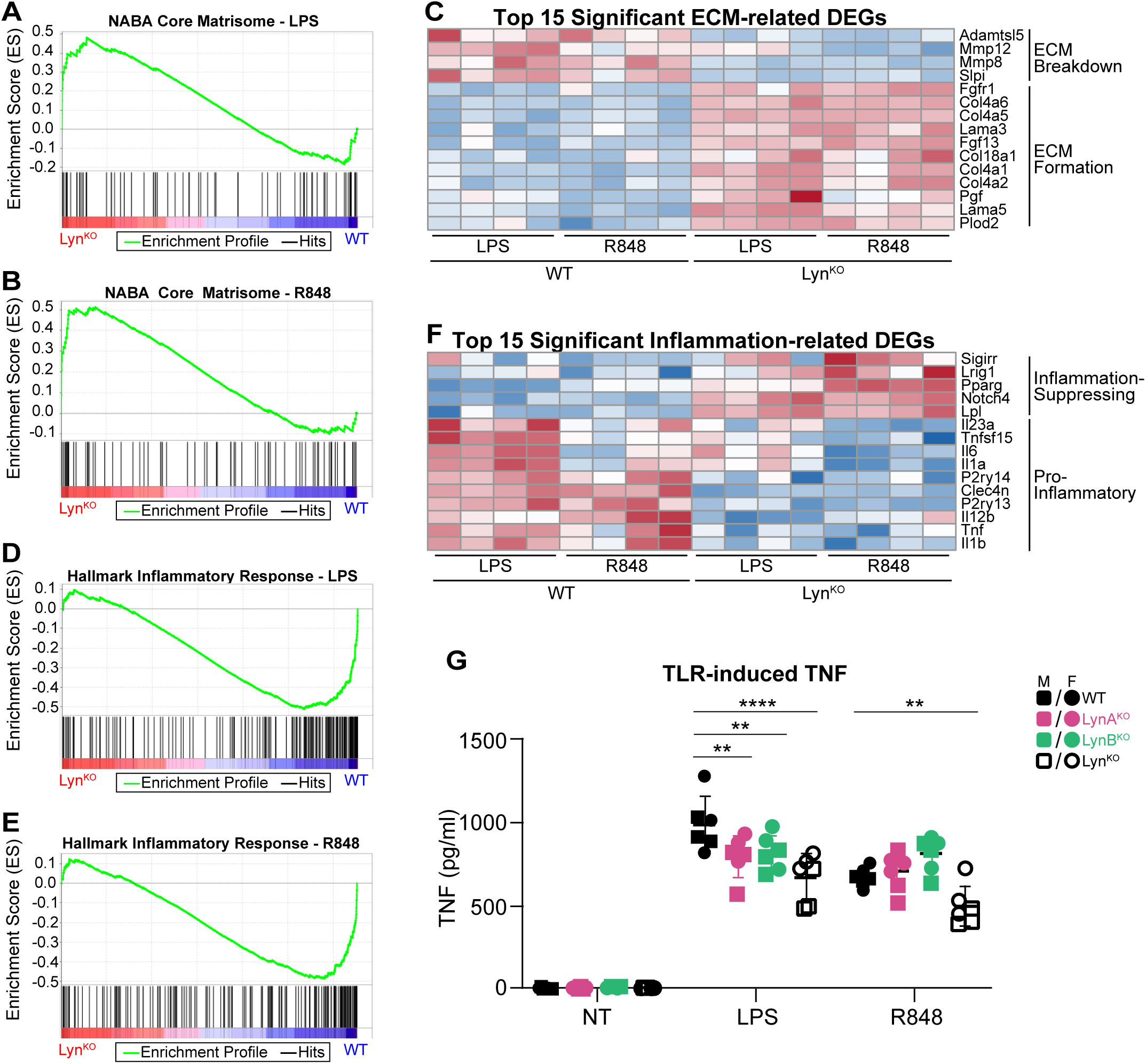
Lyn^KO^ BMDMs have enhanced expression of genes driving ECM synthesis and impaired inflammatory cytokine production after TLR stimulation. **(A,B)** GSEA showing enrichment of core matrisome genes in Lyn^KO^ BMDMs relative to WT after **(A)** LPS or **(B)** R848 treatment. **(C)** Heat map of the 15 most significant DEGs related to ECM formation in WT and Lyn^KO^ BMDMs. (D,E) GSEA showing enrichment of inflammatory response genes in WT and Lyn^KO^ BMDMs after **(D)** LPS or **(E)** R848 treatment. **(F)** Heat map of the 15 most significant DEGs related to inflammatory response in WT and Lyn^KO^ BMDMs. Heat maps and DEGs were compiled as described in Fig. 2. **(G)** ELISA showing TNF production by WT, LynA^KO^, LynB^KO^, and total Lyn^KO^ BMDMs at steady state and after 24 h treatment with 2 ng/ml LPS or 20 ng/ml R848 (*n*=6). Data are presented as mean ± standard deviation. Significance was assessed via two-way ANOVA with Tukey’s multiple comparisons test. *p*-value <0.05 *, <0.01 **, <0.001 ***, <0.0001 ****, with all other pairwise comparisons lacking significant differences. Outlier analysis was performed using unbiased ROUT with Q=1%. *n* indicates the number of biological replicates, with cells from different individual mice. Squares indicate cells derived from male mice, and circles indicate cells derived from female mice.

Lastly, GSEA more broadly confirmed the impairment of TLR-induced inflammatory responses by Lyn^KO^ BMDMs. Hallmark gene sets for inflammatory response, TNF signaling via NF-ĸB, IL-6/JAK/STAT3 signaling, and complement were all underexpressed in Lyn^KO^ cells after LPS **(Fig. 7D)** or R848 **(Fig. 7E)** treatment. Lyn^KO^ BMDMs had decreased induction of genes driving inflammatory signaling (e.g., *P2ry13, P2ry14, Clec4n*) and cytokine production (e.g., *Il1a, Il1b, Il6, Il12b, Il23a, Tnf, Tnfsf15*) in tandem with failure to downregulate expression of immunosuppressive gene products (e.g., *Lpl, Lrig1, Notch4, Pparg, Sigirr*) **(Fig. 7F)**. To ensure that differences in mRNA expression were translated to the protein level, we analyzed TLR-induced TNF secretion by BMDMs after treatment with LPS or R848. Quantifying TNF secretion via ELISA, we found that Lyn^KO^ BMDMs had diminished TLR responses, secreting 2-fold less TNF protein than WT cells after treatment with LPS or R848 **(Fig. 7G)**. Although there was no isoform-specific contribution to TNF production, it seems that TLR4 and TLR7 require different total amounts of Lyn expression to function at a WT level— LPS-treated LynA^KO^ and LynB^KO^ BMDMs had impaired TNF secretion, albeit to a lesser degree than Lyn^KO^, whereas the single-isoform Lyn knockouts had no defect in R848-induced TNF production. We therefore conclude that TLR4 requires higher levels of Lyn expression than TLR7 to maintain WT-like levels of signaling.

## Discussion

In this study, we show that either LynA or LynB is sufficient to promote TLR sensitivity, matrix remodeling, and inflammatory signaling and that complete loss of Lyn disrupts these essential macrophage functions. Both at steady state and after TLR stimulation, the expression of either Lyn isoform restores most of the widespread transcriptomic changes seen in Lyn-deficient macrophages. At steady state, Lyn is responsible for restricting the expression of genes driving DNA synthesis and replication, mitosis, and cellular growth, which translates to inhibition of macrophage hyperproliferation in culture. Interestingly, despite restoring normal expression of proliferation-related genes, single-isoform expression of Lyn is ineffective at preventing macrophage hyperproliferation. Lyn also exerts transcriptional control over ECM remodeling by driving the expression of genes that promote ECM degradation and restricting genes that direct the synthesis of structural proteins and ECM components, both at steady state and after TLR activation. Lastly, Lyn plays an important role in balancing inflammatory and immunosuppressive signaling pathways downstream of TLRs. Single-isoform expression of Lyn is sufficient for normal TLR7-driven cytokine production, while TLR4-induced TNF production may require full expression of total Lyn. Regardless, macrophages do not have an isoform-specific requirement for driving TLR-induced cytokine responses. Notably, Lyn does not have a significant impact on TLR expression in macrophages, neither transcriptionally nor posttranslationally. These findings support the conclusion that expression of either Lyn isoform is sufficient to maintain most of the canonical TLR responses and suppress dysregulated ECM formation in macrophages, although inadequate expression of total Lyn may be insufficient to fully restore proliferation control.

Transcriptomic enrichment of E2F targets and mitotic spindle components in Lyn^KO^ cells supports a model in which Lyn deficiency relieves molecular checks on cell-cycle progression. This is consistent with previous reports in DCs (Scapini et al., 2009), myeloid progenitors (Harder et al., 2001), and patrolling monocytes (Roberts et al., 2020), where Lyn deficiency promotes cell proliferation and survival. The observation that both LynA^KO^ and LynB^KO^ BMDMs proliferate more than WT, despite lacking robust transcriptional activation of the same cell-cycle programs, suggests that Lyn may restrain proliferation in a dose-dependent rather than isoform-specific manner. Furthermore, the marginal increase in expression of proliferation-associated genes that is seen with a single-isoform deficiency of Lyn may be sufficient to drive a hyperproliferative response to M-CSF. These findings raise the possibility that Lyn contributes to the maintenance of macrophage quiescence under homeostatic conditions and that loss of Lyn expression tips the balance toward expansion, even in the absence of strong mitogenic cues. Given the importance of controlled macrophage turnover in resolving inflammation and maintaining tissue integrity (Gautier et al., 2013), Lyn may serve as a key regulator of macrophage population dynamics in both steady-state and inflammatory settings.

Our study also suggests that Lyn plays an underappreciated role in controlling ECM dynamics in macrophages. Lyn^KO^ BMDMs exhibit increased expression of genes encoding collagen IV, laminins, and ECM cross-linking enzymes, coupled with reduced expression of genes encoding matrix-degrading metalloproteases such as MMP8 and MMP12. This shift toward an ECM-producing, ECM-preserving phenotype could impair immune-cell trafficking and tissue remodeling, contributing to pathological fibrosis. Additional protein-level and functional studies will be needed to determine whether Lyn directly controls macrophage-derived ECM deposition and whether isoform-specific expression of LynA or LynB is sufficient. Nevertheless, these transcriptomic findings are consistent with our previous work showing increased fibrosis in aged Lyn^KO^ kidneys (Brian et al., 2022). Conversely, a macrophage phenotype that promotes ECM synthesis and limits ECM degradation may be beneficial in suppressing cancer growth and metastasis. The ECM plays a complex role in cancer progression, where increased matrix breakdown can promote cancer-cell growth and metastasis, yet a thickened ECM can impair responsiveness to chemotherapy (Henke, et al., 2020). On the other hand, a collagen-rich ECM might suppress cancer growth by limiting the availability of oxygen and nutrients (Di Martino, et al., 2022). Lyn expression in macrophages within the tumor microenvironment promotes cancer-cell growth, and Lyn-deficient macrophages delay the progression of chronic lymphocytic leukemia and prolong patient survival (Nguyen et al., 2016). Furthermore, Lyn-deficient stromal fibroblasts reduce cancer growth by acquiring a myofibroblastic phenotype, characterized by increased ECM formation and reduced production of inflammatory cytokines (vom Stein et al., 2023). Thus, treatments targeting Lyn-mediated pathways in macrophages within tumors may prove beneficial in reducing cancer growth and metastasis by reducing ECM remodeling and limiting inflammation.

The impaired inflammatory response of Lyn-deficient macrophages underscores the importance of Lyn as a positive driver of immune signaling. While several studies have shown that Lyn inhibits TLR signaling, namely in classical DCs and B cells (Tsantikos et al., 2010; Lamagna et al., 2013; Lamagna et al., 2014), our findings align with reports indicating that Lyn is required for optimal TLR-induced cytokine production in macrophages (Shio et al., 2009; Avila et al., 2012; Toubiana et al., 2015; Lim et al., 2015; Li et al., 2016; Dallari et al., 2017). This dichotomy underscores the cell-type-specific roles of Lyn, perhaps reflecting differential expression of binding partners, other negative regulators (e.g., the inositol phosphatase SHIP1), TLR adapter proteins, or SFKs, including Lyn itself. In macrophages, Lyn likely acts upstream of several pathways essential for cytokine production, including PI3K, MAPK, IRF, and NF-κB (Shio et al., 2009; Lim et al., 2015; Smolinska et al., 2008; Li et al., 2016; Toubiana et al., 2015; Avila et al., 2012). This suggests that the inflammatory phenotype observed in Lyn^KO^ mice may be driven predominantly by immune cells not of the macrophage lineage or by cell-extrinsic effects on macrophages in vivo. For instance, macrophage-related pathologies in Lyn^KO^ mice, such as glomerulonephritis, may arise from the exacerbated inflammatory environment created by dysregulated, Lyn-deficient DCs (Lamagna et al., 2013; Ma et al., 2019) and mature B cells (Tsantikos et al., 2010; Lamagna et al., 2014), rather than innate inflammatory signaling by Lyn^KO^ macrophages.

We show that either LynA or LynB can promote TLR-induced cytokine production in macrophages. Partially impaired TLR4-driven TNF production in macrophages with single-isoform Lyn expression likely results from reduced levels of total Lyn in these cells, indicating a dose-dependent rather than isoform-specific requirement for signaling. This is supported by a previous observation that even a partial loss of Lyn can promote B-cell dysregulation and autoimmunity (Tsantikos et al., 2012). Defining how Lyn modulates signaling thresholds across different myeloid subsets and downstream of different receptors will be a critical step in resolving these apparent contradictions and elucidating how Lyn orchestrates balanced immune responses.

Our findings support a model in which Lyn acts as a positive regulator of macrophage activation down-stream of TLRs, while simultaneously serving as a brake on pathological proliferation and ECM accumulation. These dual roles may reflect a broader homeostatic function for Lyn in tuning macrophage responses to inflammatory stimuli, enabling robust immune activation while limiting myeloid-cell expansion and tissue fibrosis. Given that expression of either LynA or LynB alone can restore many macrophage functions to WT-like levels, therapies aimed at boosting total Lyn expression or function could offer greater benefit than isoform-specific modulation. Future studies dissecting the mechanistic contributions of LynA and LynB to specific signaling nodes — particularly their interactions with adaptor proteins and downstream kinases — will be essential for translating these insights into therapeutic approaches.

## Authorship

AJL: Conceptualization, Methodology, Investigation, Data curation, Formal analysis, Visualization, Writing – original draft, Writing – review & editing, Funding acquisition. JTG: Methodology, Investigation, Formal analysis, Writing – review & editing, Funding acquisition. TSF: Conceptualization, Methodology, Visualization, Writing – review & editing, Supervision, Project administration, Funding acquisition.

## Acknowledgements

We thank Dr. Tyler Bold, Dr. Jesse Williams, Dr. Natalia Calixto Mancipe, Yingzheng Xu, and Charlie Roll for contributions to analysis and presentation of RNA sequencing data. Thanks also to Drs. Bryce Binstadt, Vaiva Vezys, Tyler Bold, Jesse Williams, and Li-Na Wei for feedback on project goals and presentation.

## Data Availability

RNA sequencing data have been deposited in the NCBI Gene Expression Omnibus (GEO).

## Abbreviations

AP1: Activator protein 1
BMDM: Bone-marrow-derived macrophage
CARD9: Caspase recruitment domain-containing protein 9
CCL: C-C motif chemokine ligand
cDC: Classical dendritic cell
CpG: Cytosine-phosphate-guanine dinucleotide
CREB: Cyclic AMP-responsive element-binding protein
CTV: CellTrace^TM^ violet
CXCL: C-X-C motif chemokine ligand
DC: Dendritic cell
DEG: Differentially-expressed gene
DMEM: Dulbecco’s modified Eagle medium
ECM: Extracellular matrix
ELISA: Enzyme-linked immunosorbent assay
GSEA: Gene set enrichment analysis
IFN: Interferon
IL: Interleukin
IRF: Interferon regulatory factor
KO: Knockout
LPS: Lipopolysaccharide
MAPK: Mitogen-activated protein kinase
M-CSF: Macrophage colony-stimulating factor
MMP: Matrix metalloprotease
MYD88: Myeloid differentiation primary-response protein 88
NF-ĸB: Nuclear factor kappa B
NLRP3: NOD-, LRR-, and pyrin domain-containing protein 3
PCA: Principal component analysis
pDC: Plasmacytoid dendritic cell
PI3K: Phosphoinositide-3 kinase
RIG: Retinoic acid-inducible gene
SFK: Src-family kinase
SHIP: Src homology 2 domain-containing inositol polyphosphate 5-phosphatase
TBK: TANK-binding kinase
TLR: Toll-like receptor
TNF: Tumor necrosis factor
TRIF: TIR-domain-containing adaptor inducing interferon-beta
VST: Variance-stabilizing transformation
WT: Wild type

---

**Supplemental Figure 1.**
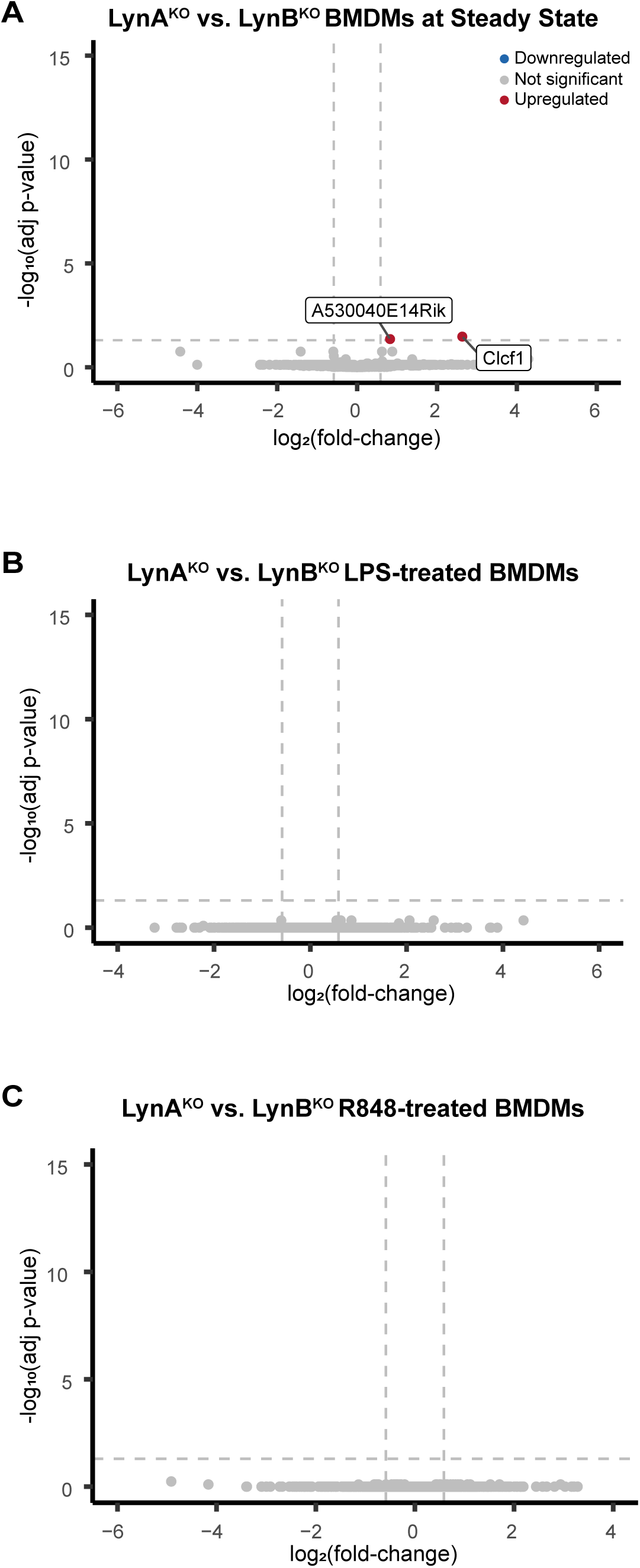
LynAKO and LynB^KO^ BMDMs have similar transcriptomes at steady state and after TLR stimulation. **(A-C)** Volcano plots highlighting DEGs between LynA^KO^ and LynB^KO^ BMDMs at **(A)** steady state and after **(B)** LPS or **(C)** R848 treatment. DEGs were calculated as described in *Fig. 2*.

**Supplemental Figure 2.**
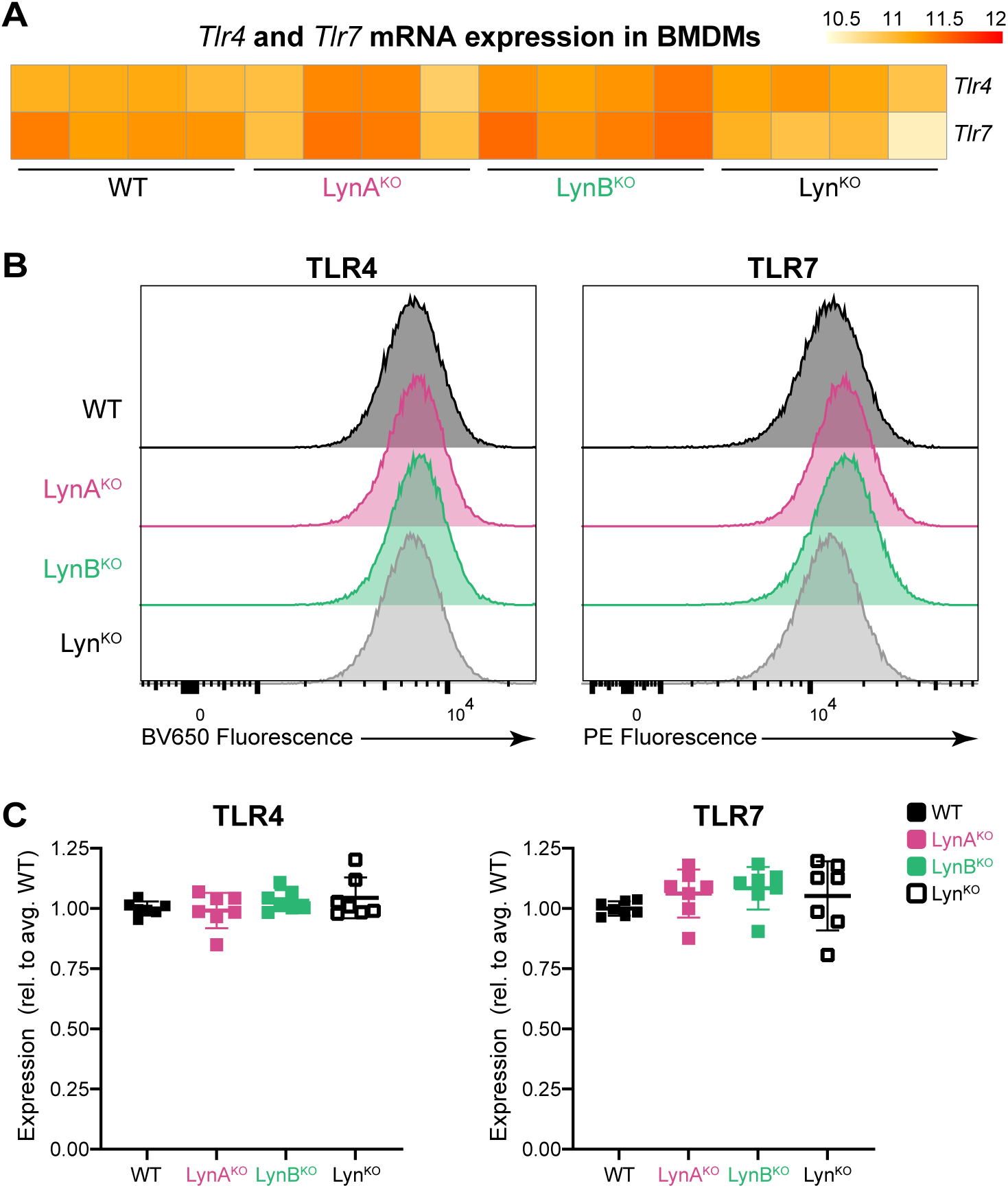
Lyn knockout does not affect TLR4 or TLR7 expression by BMDMs. **(A)** RNA-sequencing data showing VST-normalized hit-counts of *Tlr4* and *Tlr7* in BMDMs at steady state (red indicates higher expression). **(B)** Representative flow-cytometry histograms showing protein-level expression of TLR4 (BV650 anti-mouse CD284/MD-2 Complex) and TLR7 (PE anti-mouse CD287) in BMDMs. **(C)** Quantified flow cytometry data showing relative TLR expression on the surface of WT and Lyn-deficient BMDMs, comparing the geometric mean fluorescent intensity (gMFI) for each sample to the gMFI of WT within each experiment (*n*=7 biological replicates over 3 experimental days). No significant differences were observed.

**Supplemental Figure 3.**
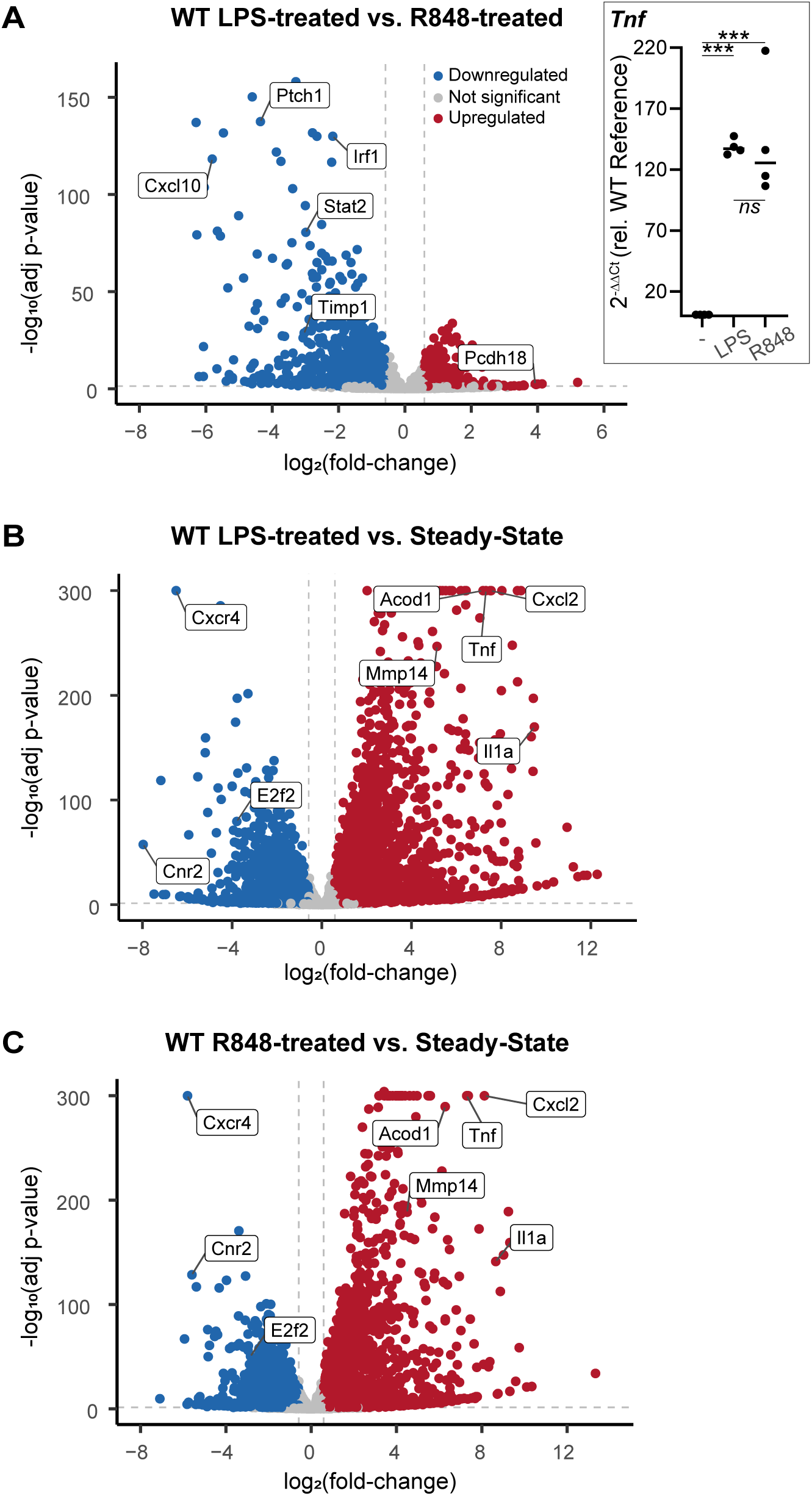
LPS and R848 have differential impacts on gene transcription in WT BMDMs. **(A inset)** qRT-PCR analysis of *Tnf* expression in response to 2 h treatment with 2 ng/ml LPS or 20 ng/ml R848. Significance was assessed via one-way ANOVA with Tukey’s multiple comparisons test: \*\*\**P* = 0.0002-0.0003. There was no significant difference (*ns*) between LPS and R848 conditions (*P*=0.9669). **(A-C)** Volcano plots highlighting DEGs in **(A)** LPS- or R848-treated WT BMDMs relative to each other, **(B)** LPS-treated relative to steady-state, or **(C)** R848-treated relative to steady-state. DEGs were calculated as described in *Fig. 2*.

**Supplemental Figure 4.**
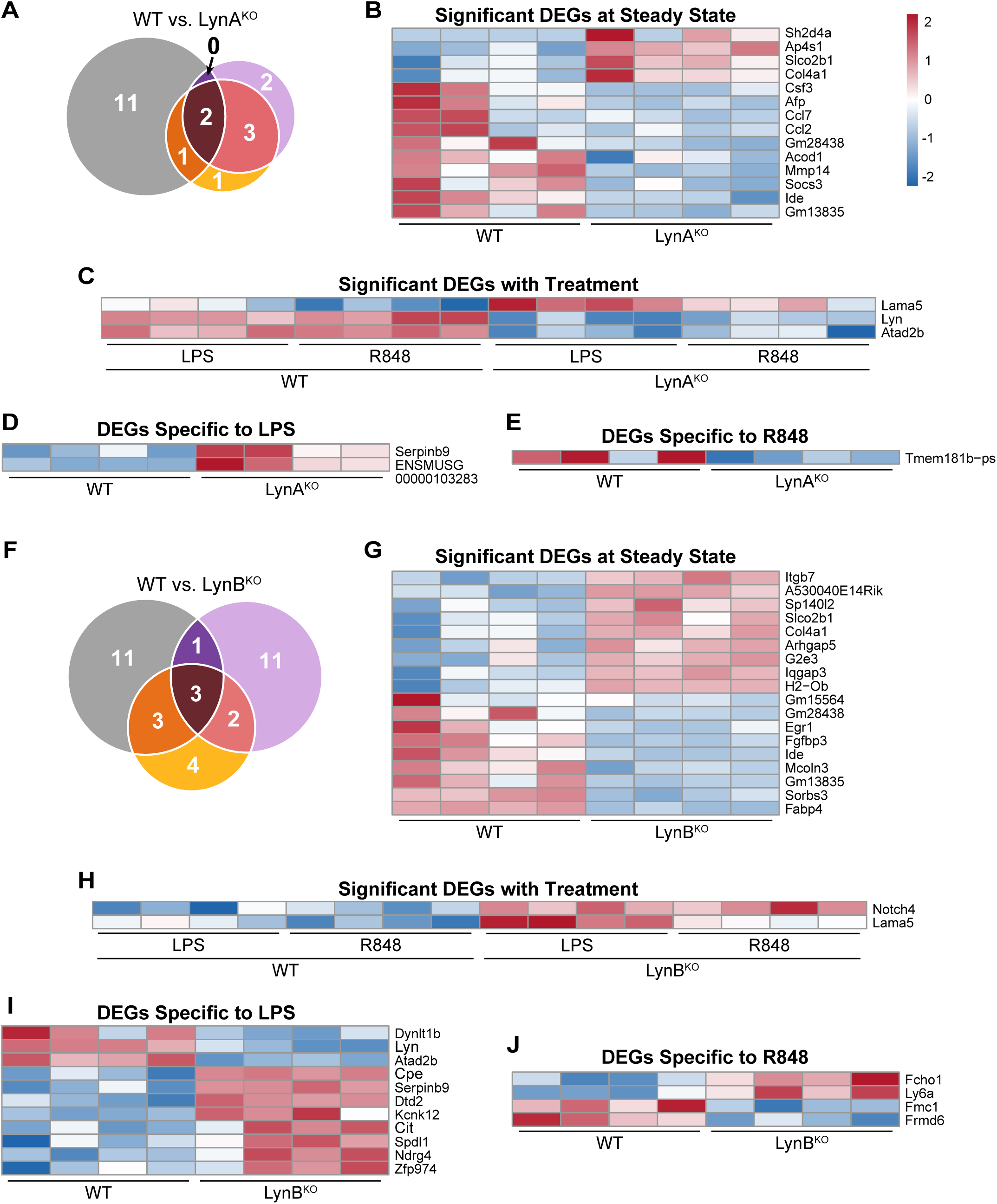
LynAKO and LynB^KO^ BMDMs have few transcriptomic differences with WT cells. **(A,F)** Venn diagrams illustrating significant DEGs between WT and **(A)** LynA^KO^ or **(F)** LynB^KO^ BMDMs across all treatments. **(B-J)** Heat maps of all significant DEGs between WT and **(B-E)** LynA^KO^ or **(G-J)** LynB^KO^ BMDMs at **(B,G)** steady state, **(C,H)** after either LPS or R848 treatment, **(D,I)** only after LPS treatment, and **(E,J)** only after R848 treatment. Heat maps and DEGs were compiled as described in *Fig. 2*.

**Supplemental Figure 5.**
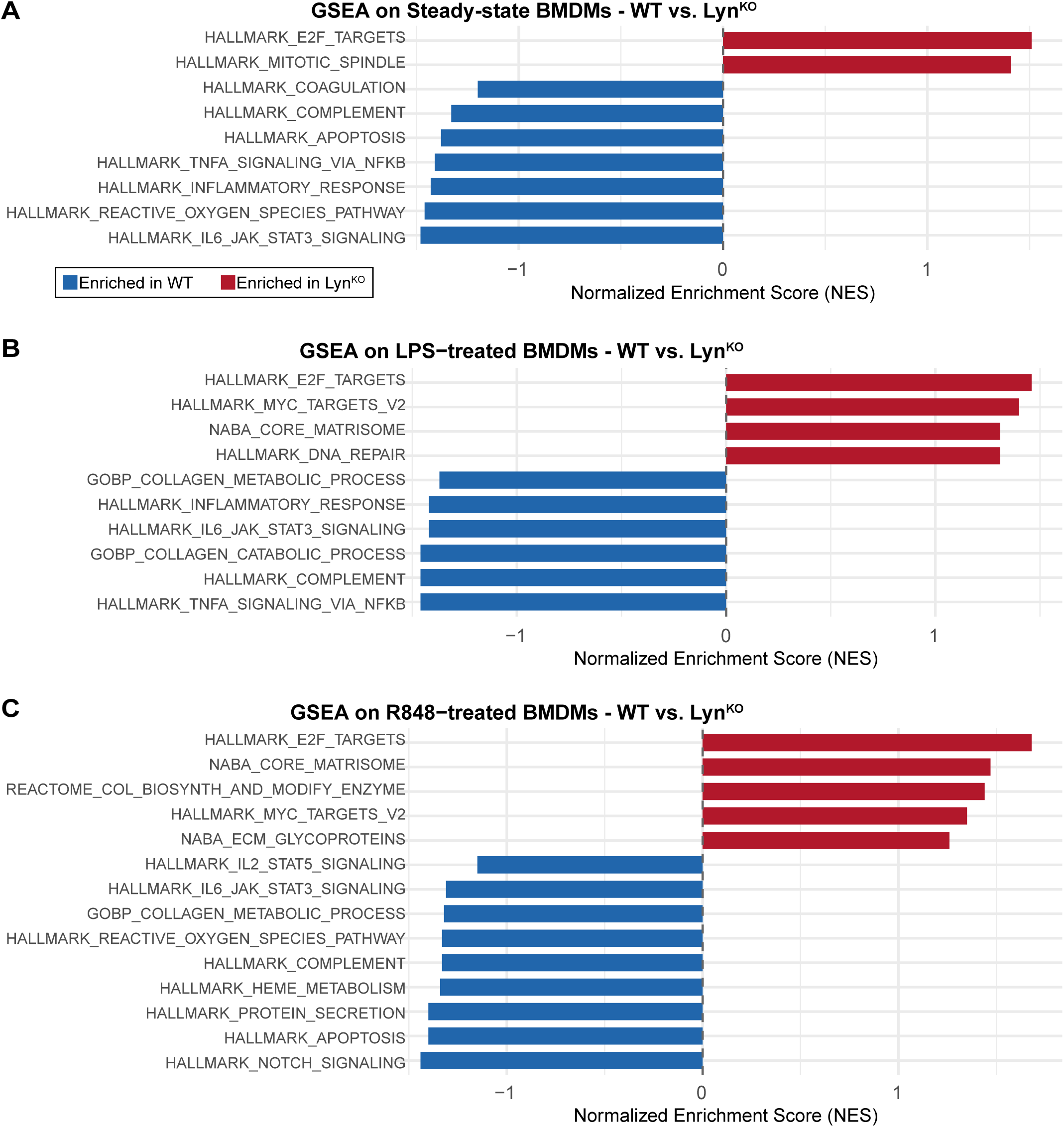
Enrichment of cell-cycle and matrix-assembly pathways in Lyn^KO^ BMDMs; enrichment of inflammatory and catabolic pathways in WT. Graphical summary of GSEA performed on RNA-sequencing data from WT and Lyn^KO^ BMDMs at **(A)** steady state and after **(B)** LPS or **(C)** R848 treatment (*n*=4). Bar plots show normalized enrichment scores (NES) for significantly enriched pathways identified using GSEA, with hallmark and curated gene sets from the MSigDB. Positive NES values (red) indicate enrichment in Lyn^KO^; negative NES values (blue) indicate enrichment in WT. Significance was defined by a nominal *p*-value <0.1. *n* indicates the number of biological replicates per genotype (each from a different mouse) and treatment.

